# Telomeres as integrative markers of exposure to stress and adversity: A systematic review and meta-analysis

**DOI:** 10.1101/320150

**Authors:** Gillian V. Pepper, Melissa Bateson, Daniel Nettle

## Abstract

Telomeres have been proposed as a biomarker that integrates the impacts of different kinds of stress and adversity into a common currency. There has as yet been no overall comparison of how different classes of exposure associate with telomeres. We present a meta-analysis of the literature relating telomere measures to stresses and adversities in humans. The analysed dataset contained 543 associations from 138 studies involving 402,116 people. Overall, there was a weak association between telomere variables and exposures (greater adversity, shorter telomeres: *r* = −0.15, 95% CI - 0.18 to −0.11). This was not driven by any one type of exposure, since significant associations were found separately for physical diseases, environmental hazards, nutrition, psychiatric illness, smoking, physical activity, psychosocial and socioeconomic exposures. Methodological features of the studies did not explain any substantial proportion of the heterogeneity in association strength. There was, however, evidence consistent with publication bias, with unexpectedly strong negative associations reported by studies with small samples. Restricting analysis to sample sizes greater than 100 attenuated the overall association substantially (*r* = −0.09, 95% CI −0.13 to −0.05). Most studies were underpowered to detect the typical association magnitude. The literature is dominated by cross-sectional and correlational studies which makes causal interpretation problematic.

## Introduction

Exposure to stress and adversity across the lifespan is associated with increased morbidity and mortality from many causes. This implies that stress and adversity have a lasting impact on general physiological processes ‘under the skin’. However, until recently, there were few candidate markers of this accumulation of physiological damage. In the last 15 years, the idea that telomeres might serve such a role has rapidly gained in scientific popularity. In particular, telomeres offer a potential ‘psychobiomarker’ that integrates the organism’s experience of psychological states, social and environmental contexts, as well as physical damage, into a common currency [1]. Telomeres are DNA-protein complexes that form protective caps on the ends of chromosomes, and are thought to play a key role in preserving chromosomal stability. At the cellular level, critically short telomere length leads to replicative senescence. At the whole organism level, average telomere length reduces with age. Thus, telomere length or attrition is a biomarker of ageing. Since the impact of stress and adversity may be to increase the individual’s biological age (as opposed to chronological age), telomere measures offer a metric with which to assess Hans Selye’s famous contention that: ‘Every stress leaves an indelible scar, and the organism pays for its survival after a stressful situation by becoming a little older’ [2].

Interest in using telomeres as a ‘psychobiomarker’ has grown rapidly, not just in human epidemiology, but also in animal ecology [3], and animal welfare [4]. In the human literature, telomeres have been studied in association with a wide range of exposure variables, including psychological stress [5], psychiatric illness [6], socioeconomic status [7], environmental pollutants [8], nutrition [9], smoking [10] and physical activity [11]. In several of these cases, the number of studies is sufficient that meta-analyses have appeared [12–20], often finding that telomere length is associated with the exposure, though weakly and variably. Reviewing the associations between telomeres and different exposures separately is appropriate to answer questions about that particular exposure. However, it loses sight of the most exciting promise of telomeres as a ‘psychobiomarker’: namely their potential to integrate the consequences of quite different kinds of stress and adversity into a common currency. Here, we set out to simultaneously review relationships of telomere length and attrition with all the *different* kinds of stress and adversity that are being studied.

Having a single integrated dataset allows several possibilities not available in separate, specialist meta-analyses. First, it offers a synopsis of the whole burgeoning field of telomere epidemiology. Second, it allows explicit comparison of different association strengths on the same scale (is the association of telomeres with psychological stress generally weaker or stronger than the association with physical disease, or exposure to pollution?). Third, it offers potential to address unanswered questions about telomere dynamics, such as whether, overall, early-life stressors are more strongly associated with telomere shortening than stressors experienced in adult life, as has been suggested [21]. Fourth, it leads to a large dataset within which some methodological issues of broad relevance can be examined. These include whether different tissue types produce different patterns, and whether the popular telomere-length measurement method, quantitative polymerase chain reaction (qPCR) [22], leads to generally weaker associations than other methods. This should be expected, since measurement error is generally found to be higher in qPCR than more intensive methods [23], and measurement error attenuates observed correlations.

With these objectives in mind, we carried out a systematic review and meta-analysis of the published literature on telomeres in relation to stress and adversity *in vivo*, to May 2016. Our searches involved the terms ‘stress’ and ‘adversity’ in combination with ‘telomere’. We extracted all associations with telomeres reported in the papers, not just the associations of primary interest to the original authors. Our search strategy was not intended to find the whole of the literature on telomeres and any particular exposure variable. Authors may not always have used the descriptors we searched, and may have been more likely to do so for some exposure variables than others. Nonetheless, our searches did produce the largest telomere literature dataset assembled to date, and we believe that though not exhaustive, it constitutes a good transect through the field of telomere epidemiology.

## Methods

Our methods are described in detail in our protocol, which was registered via the Open Science Framework prior to data extraction [24]. Raw data files and data analysis scripts are freely available in an online archive on the Zenodo repository [25].

### Search strategy and inclusion criteria

A PRISMA diagram for our study is available as electronic supporting material. We searched the Scopus and PubMed databases for papers including the words “stress” or “adversity”, and “telomere”. All records up to the date of the search (11^th^ May 2016) were screened (n = 3647). We removed duplicates and t hen screened the remaining papers based on their titles and abstracts. In summary, this involved removing any papers that: 1) were not complete original research papers available electronically and in the English language; 2) used study organisms from outside the animal kingdom; 3) did not study whole organisms; 4) used genetically modified organisms; 5) experimentally applied non-naturalistic exposures in captive animals; 6) examined telomere length in transplanted tissues or organs; 7) were presented as concerning the physiological consequences of telomere length, rather than the correlates of exposures; 8) examined intergenerational questions (e.g. the effects of paternal infection status on offspring telomere length); or 9) used the same data set, or participants reported in a previously recorded paper, to address a exposure-telomere relationship we had already recorded (where this occurred, the first-recorded association was the one used). This led to a candidate set of 286 papers.

Although our searches were based on the terms “stress” and “adversity”, we extracted all reported associations with potential exposure variables found in the papers returned by our search, whether or not they were the focus of the study’s stated objectives. This included control variables and covariates as long as sufficient detail was provided. Thus, our search strategy consisted of finding papers on the subjects of stress/adversity and telomeres, and sampling the full variety of exposures that fell out of the papers captured by the search.

### Association format

Associations could only be used if convertible into a correlation coefficient, the common association metric that we chose based on initial scoping and piloting. Standard conversion formulae were used [26–29], and the conversion algorithms are provided in the data repository. Usable statistics comprised: correlation coefficients, standardised βs from regression models; unstandardized βs where standard deviations for the independent and dependent variables were provided; unstandardized betas from regressions with dichotomous independent variables where the standard deviation of the dependent variable was provided; F-ratios from ANOVAs comparing two groups; t-statistics; Cohen’s *d* or standardised mean difference statistics; group means with standard errors; and group means with standard deviations. Where several alternative analyses were presented, we chose unadjusted in preference to adjusted analyses; and from several adjusted analyses where no unadjusted data were available, we chose the analysis that adjusted for the fewest variables. This was to maximise comparability between studies that used different sets of control variables in multivariate models. For longitudinal studies, associations were between change in telomere length (i.e. difference between follow-up and baseline) and the exposure variable. For 125 papers, the reported information was not sufficient to create a usable correlation coefficient.

### Data extraction

Data from 218 associations (30%) were extracted independently by both GP and DN. Any differences were identified and resolved as part of the process of refining our data extraction methods. The remaining associations were extracted by either GP or DN, with GP checking and correcting the whole dataset after extraction. As well as sample size and other statistics necessary for association conversion, we extracted bibliographic information and a series of classificatory variables as described in table 1. The life stage prior to birth was classed as embryonic, and the life stage prior to sexual maturity (4745 days for human females / 5,110 days for males [30]) was defined as childhood. We also identified any associations that could be considered sub-parts of others, for example, separate-by-sex associations where the combined association was also reported, or associations between telomeres and sub-scales where the association with the main scale was also reported. We did not include these sub-scale and sub-group associations in our final analyses, though they are included in the unprocessed data set, in case they are of interest to others.

**Table 1.**
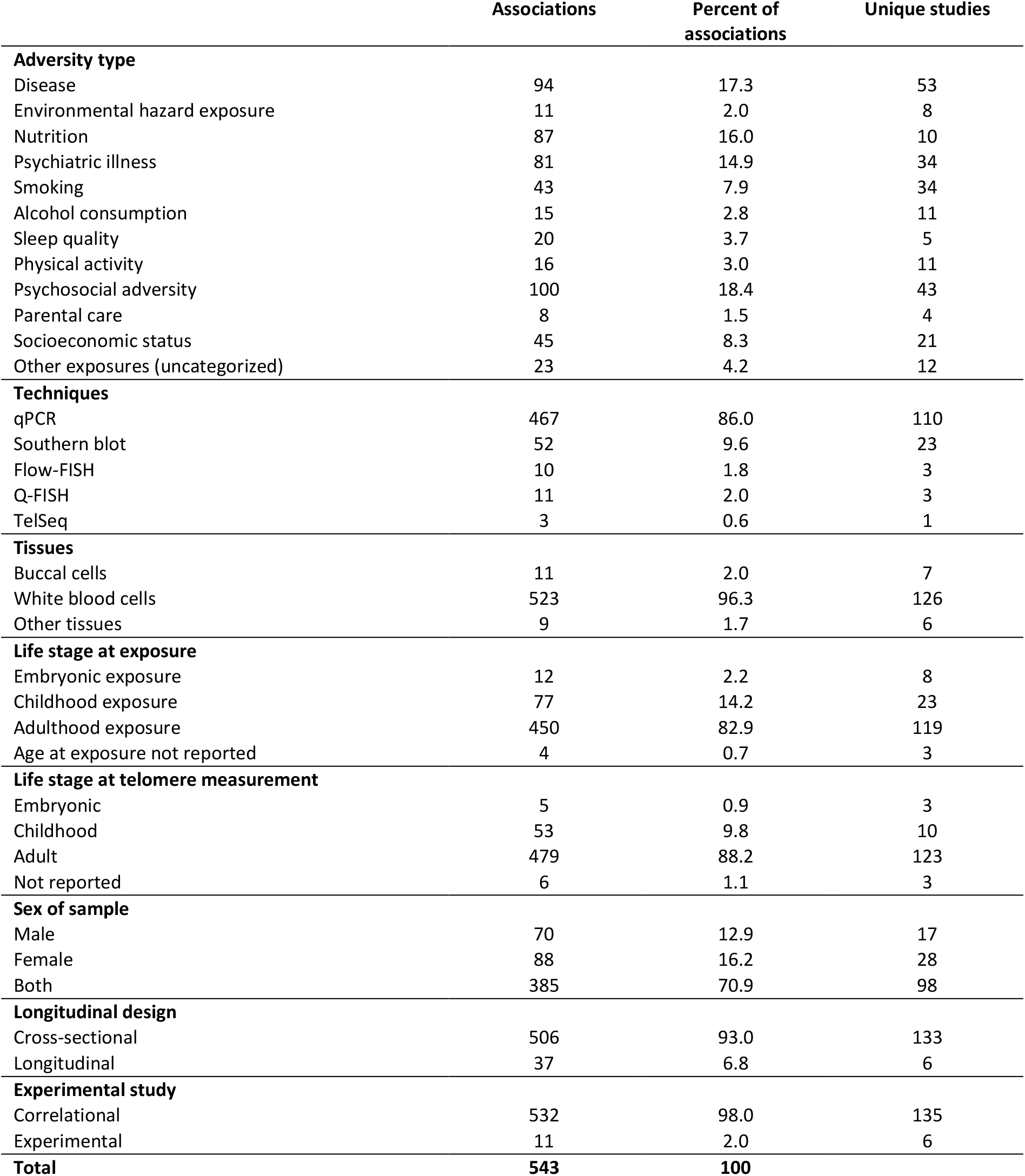
Characteristics of the associations included in the analysis. The numbers of unique studies for each category do not sum to the number of studies in the whole dataset (138), as some studies contribute associations in several categories.

### Final dataset

The extracted data are the ‘unprocessed data’ file in the data archive. We categorized exposure variables *a posteriori*. We created both broad categories (11 categories plus ‘Other’, as specified in table 1), and fine ones, for example using specific diseases rather than ‘physical disease’, and specific types of psychosocial or socioeconomic measures (35 fine categories, plus ‘Other’). No categories (broad or fine) were created if the number of independent associations (i.e. from different papers) was less than 3 prior to the exclusions described below.

The final dataset analysed here (‘processed data’ file) differs from the unprocessed data in a number of regards. We excluded 27 associations from studies of non-human animals, since we deemed these too few (typically one study per species) for further analysis. We then excluded 11 associations that appeared to be duplicates of those reported in other papers using the same data, and 14 associations where the exposure variable was a medical treatment—the designs of these studies generally confounded effects of the treatment on telomeres with effects of the disease the treatment was for. Finally, we excluded 129 associations based on subscales and sub-groups of other associations reported in the same studies.

For analysis, we reversed 149 correlations in sign to align all correlations into the same direction; that is, so that a negative correlation indicates that greater stress and adversity is associated with short (or shortening), rather than long (or lengthening), telomeres. For example, we reversed correlations where the exposure variable represented higher socioeconomic status, better sleep, or more parental care, to align them with the more common case where a higher value of the exposure indicates more adversity (e.g. disease, psychosocial stress, pollution exposure). The case of nutritional variables was challenging, since it was often unclear if higher consumption was predicted to be positive or negative in effect. We therefore reversed the direction of all nutritional variables, so that a negative correlation means a deficit in consumption is associated with short telomeres. This had little impact on the results, since the overall associations for many nutritional categories were null. The categories with the strongest associations—fruit, legumes and vegetables, and vitamins—are cases that clearly conform to our assumed ‘more is better’ principle.

### Data analysis

Data were meta-analysed in R [31] using the ‘metafor’ package [32]. Estimation was by REML. Since the dataset includes multiple associations from the same studies, we used multilevel models containing nested random effects of association and study. Meta-regression was used to examine differences in association strength for different types of exposure, and different methodological features. Additional analyses to detect outliers and account for possible publication bias patterns were implemented using R packages ‘metaplus’ [33] and ‘weightr’ [34] and are reported in a separate supplementary analysis appendix. The full data analysis script is included in the data archive.

## Results

### Description of dataset

The final data set consisted of 543 associations from 138 unique studies of human participants (associations per study 1-43, mean 3.93). One hundred and sixty-eight associations were reported by the study authors as being statistically significant, 349 as null, and 26 were not reported as either null or significant. Two hundred and ninety-four associations (54%) were completely unadjusted; the remaining 249 (46%) featured some degree of statistical adjustment (for example, for age, though the exact specification of the adjustment varied from study to study). Table 1 describes the associations and studies included. Typically, they used qPCR to measure telomeres; did so in leucocytes or whole blood; were correlational rather than experimental; and were cross-sectional rather than longitudinal. This meant that the telomere variable was overwhelmingly a single measure of average length, rather than the rate of attrition. Associations with both length and attrition are included in our main analyses, though we test whether study design moderates observed correlations. The studies were mostly of adults, and mostly related to sources of stress and adversity that were experienced in adulthood.

### Overall association and publication bias

The distribution of correlations between exposure measures and telomeres is shown in figure 1a. Though the modal correlation was close to zero, the distribution was asymmetric, with 399 correlations less than zero (indicating that greater adversity was associated with shorter telomeres), and 144 greater than zero (indicating that greater adversity was associated with longer telomeres). The majority of correlations fell into what is conventionally defined as a ‘small’ effect size (−0.2 < *r* < 0.2), with 294 small negative correlations and 131 small positive ones. There were 105 instances of a moderate or large negative association, against just 13 of a moderate or large positive association. In a simple meta-analytic model with no moderators, the overall estimate of association was conventionally small and significantly negative (*r* = −0.15, 95% CI −0.18 to −0.11, p<0.001). Thus, greater adversity was correlated with shorter telomeres. There was substantial heterogeneity between associations (τ = 0.21, Q_542_ = 12742.54, p<0.001), and most of the variability resided at the between-study level rather than between associations from the same study (ρ represents the intraclass correlation coefficient between the associations from the same study; ρ = 0.87). In the supplementary analysis document (section 2), we present evidence of over-dispersion of associations relative to a Normal distribution: twelve studies were classed as outliers, ten reporting very strong negative associations, and two reporting moderately strong positive associations.

**Figure 1.**
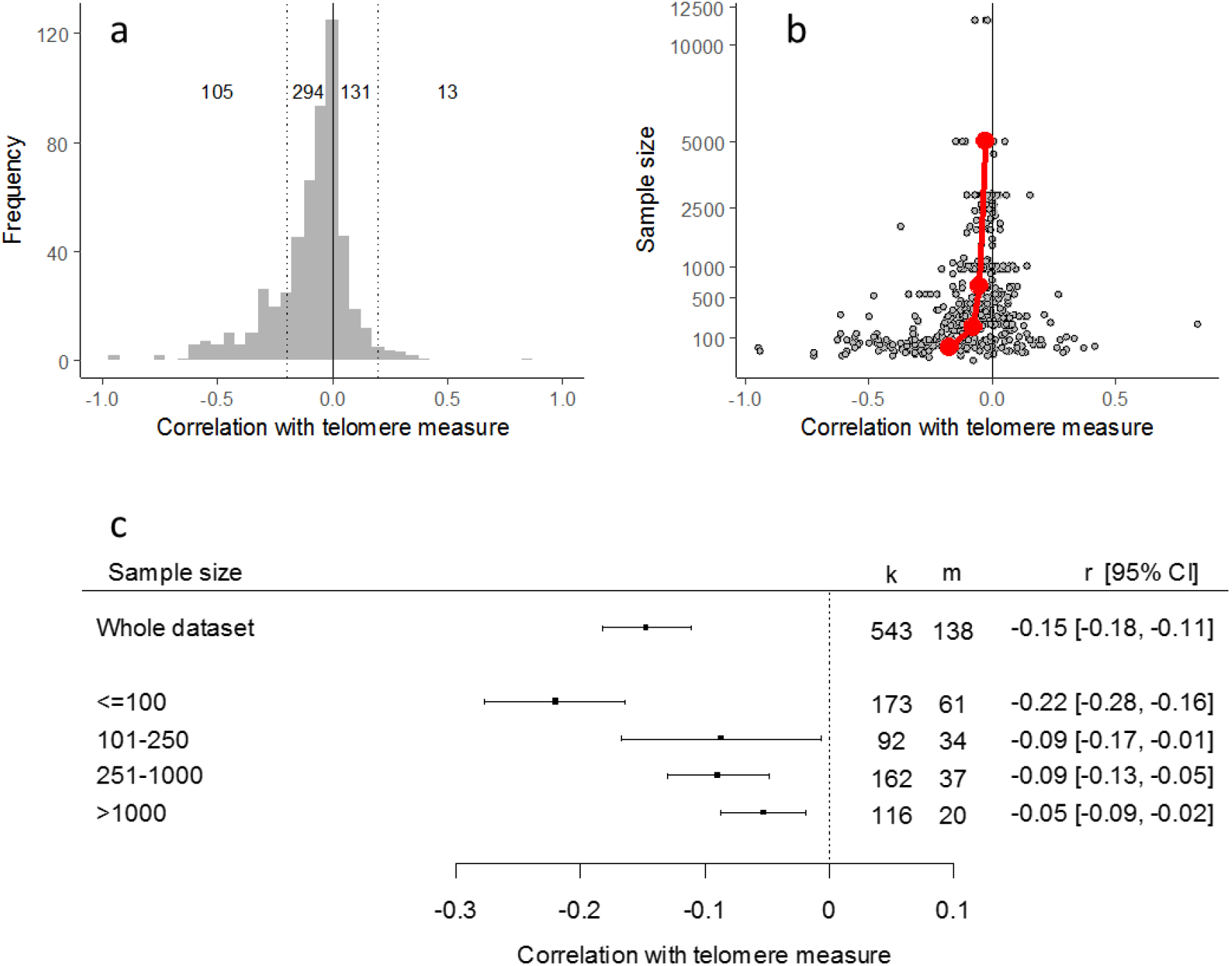
Features of the meta-analysed data. a. The distribution of correlations between exposures and telomere length or telomere attrition in the 543 associations of the whole dataset. The numbers give the number of correlations in the moderate or strong negative (*r <=* −0.2), small negative (−0.2 < *r <* 0), small positive (0 < *r <* 0.2), and moderate or strong positive (*r >*= 0.2), effect size bins respectively. b. Funnel plot of sample size against observed correlation between telomere measure and exposure variable. Red points represent the mean correlation observed for sample sizes in bins fewer than 100; 101-250; 251-1000; and more than 1000. c. Forest plot of central correlation estimate and 95% confidence interval for the whole dataset, and separately for the four bins of sample size. The *k* column represents the numbers of correlations and the *m* column the number of unique studies.

The estimated association may have been affected by publication bias. The funnel plot of correlation coefficient against sample size showed the inevitably broader range of observed correlations at smaller sample size. However, the funnel was asymmetric, with strongly negative correlations appearing at small sample sizes, but rather few of the strongly positive correlations that ought also to be expected, given that true effects were weak and the precision of estimation low in small studies (figure 1b; see also supplementary analysis document, section 3). To further explore this, we divided sample sizes into four bins (fewer than 100, 101-250, 251-1000, more than 1000; these bins represent approximate quartiles of sample size). We then added sample size bin to the meta-analytic model as a moderator (this is the conceptual equivalent of the Egger test for the multilevel model situation). Sample size bin explained a significant amount of variability (Q_3_ = 30.27, p<0.001), though substantial heterogeneity remained (τ = 0.19). In particular, associations with sample sizes of ‘fewer than 100’ were significantly more negative than the reference category of more than 1000 (B = − 0.17, 95% CI −0.24 to −0.10, p < 0.001; figure 1c; see also supplementary analysis document, section 2). The other sample size bins did not differ significantly from the reference category of ‘more than 1000’. Because the trim and fill methodology for imputing the associations assumed to be missing is not defined for the multilevel situation, we performed all subsequent analyses both on the whole dataset and, in parallel, on only the 370 correlations from 82 studies where the sample size was greater than 100 (henceforth the ‘reduced dataset’). The central estimate from the reduced dataset was considerably weaker than the full dataset (*r* = −0.09 compared to *r* = −0.15, 95% CI −0.13 to −0.05, p<0.001), with substantial heterogeneity (τ = 0.18, Q_369_ = 4512.37, p < 0.001), and again, most of the variation residing between studies, rather than between associations within studies (ρ = 0.88).

### Categories of exposure

We divided our exposure variables into 11 broad categories plus ‘other’ and added category of exposure to the meta-analytic model as a moderator. Exposure category did not explain a significant amount of the heterogeneity (whole dataset: Q_11_ = 17.25, p = 0.10; reduced dataset: Q_11_ = 14.70, p = 0.20), suggesting that the type of exposure studied, at this coarse level, does not explain the variation in association between telomeres and exposure variables. We also fitted separate meta-analytic models to the correlations in each of the 12 broad exposure categories (figure 2). In the whole dataset, the central estimate of association was numerically negative for all categories, and significantly so for all except alcohol, sleep, parental care and ‘other’. In the reduced dataset, environmental hazard additionally became non-significant. In some categories, excluding small studies markedly reduced the central correlation estimate, for example: psychosocial (from *r* = −0.16 to *r* = −0.06); psychiatric illness (from *r* = −0.13 to *r* = −0.08); and physical disease (from *r* = −0.15 to *r* = −0.11). In other categories, such as smoking, socioeconomic status, and physical activity, excluding the smaller studies had minimal effect on the (already weaker) central correlation estimates.

**Figure 2.**
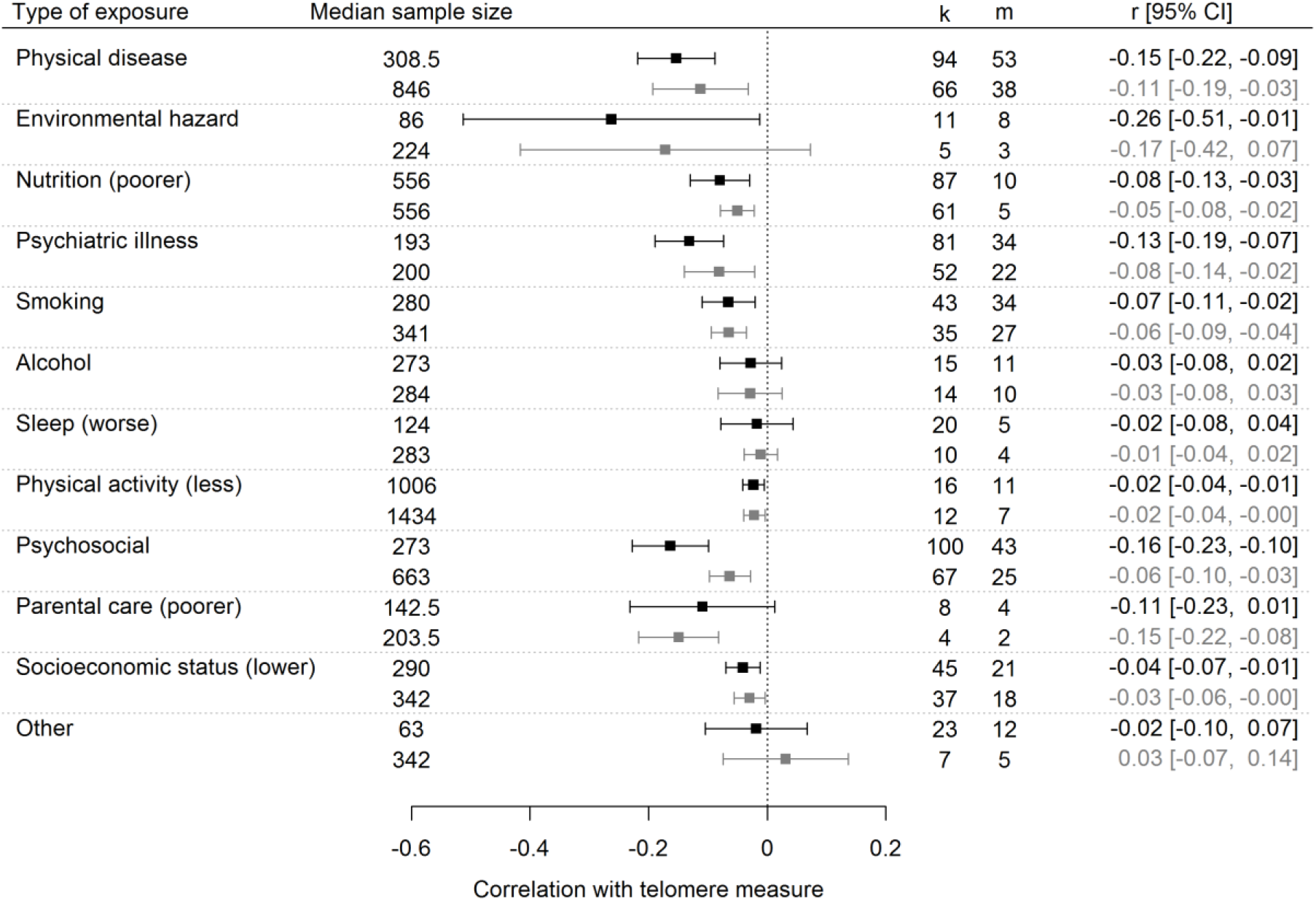
Forest plot showing central estimates of correlation and 95% confidence intervals from meta-analytic models for each broad category of exposure separately. For each category, the first row represents the full dataset, and the second, the reduced dataset (only sample sizes of 100 or greater). The *k* column represents the numbers of correlations and the *m* column the number of unique studies.

We also created a finer 36-category classification of exposures. For example, we considered each physical disease, psychiatric condition or psychosocial construct for which multiple independent data points were available separately. The fine categories explained a significant amount of heterogeneity in both the full (Q_35_ = 59.28, p < 0.01; τ = 0.20) and reduced datasets (Q_32_ = 58.03, p < 0.01; τ = 0.16; note only 33 of the 36 fine categories were represented in the reduced dataset). We took smoking as the reference category as this association is estimated with good precision due to a large number of studies. Compared to smoking, in the full dataset, we found significantly stronger negative correlations for environmental hazards (B = −0.15, 95% CI −0.26 to −0.05, p < 0.01); HIV & AIDS (B = −0.15, 95% CI −0.28 to −0.02, p = 0.02); schizophrenia (B = −0.18, 95% CI −0.34 to −0.01, p = 0.04); and lower vitamin consumption (B = −0.22, 95% CI −0.37 to −0.07, p < 0.01). Parkinson’s disease gave a significantly weaker negative correlation than smoking (B = 0.26, 95% CI 0.06 to 0.45, p < 0.01). In the reduced dataset, the significant differences from smoking for environmental hazards and schizophrenia became non-significant; the significant differences persisted for HIV & AIDS (B = - 0.14, 95% CI −0.25 to −0.03, p = 0.02) and Parkinson’s (B = 0.41, 95% CI 0.21 to 0.60, p < 0.001); and two new significant differences were found: poor parental care gave significantly stronger negative correlations than smoking (B = −0.14, 95% CI −0.27 to −0.01, p = 0.04), and low carbohydrate consumption gave significantly weaker ones (B = 0.09, 95% CI 0.00 to 0.19, p = 0.04). We also considered whether the associations in the 36 fine categories differed significantly from zero when considered separately (figure 3). In the full dataset, anxiety, cardiovascular disease, depression, diabetes, lower education, environmental hazards, lower fruit, legume and vegetable consumption, lower income, lower meat, fish and egg consumption, lower physical activity, PTSD, schizophrenia, smoking, stress, traumatic experience, and lower vitamin consumption were all significantly correlated with shorter telomeres. Restricting consideration to the reduced dataset, the associations remaining significantly less than zero were: anxiety; cardiovascular disease; diabetes; education; lower fruit, legume and vegetable consumption; lower meat, fish and egg consumption; lower physical activity; and smoking. In addition, the parental care correlation (receiving poorer care associated with shorter telomeres), which had not been significantly different from zero in the full dataset, became so in the reduced dataset.

**Figure 3.**
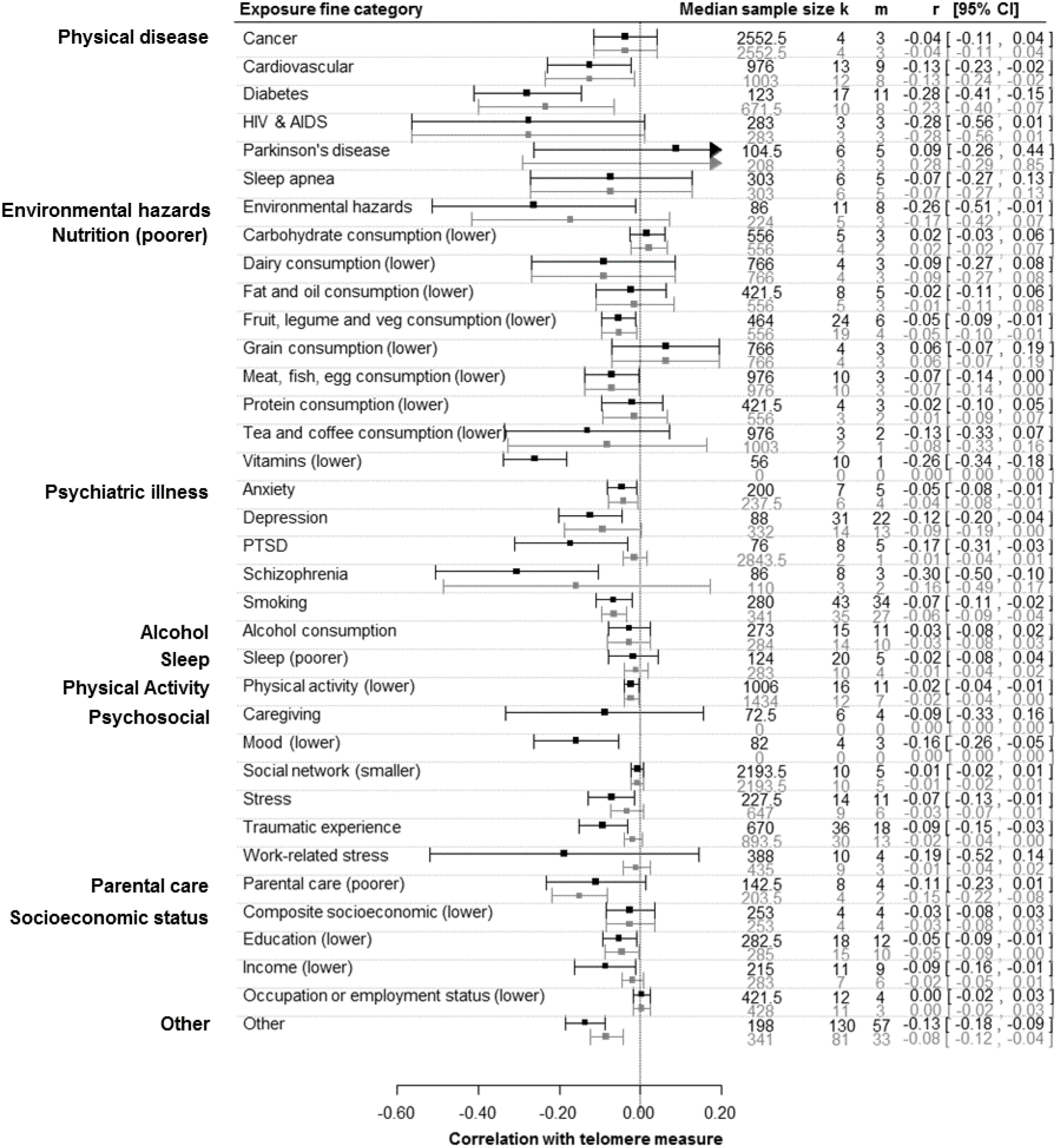
Central estimates of correlation and 95% confidence intervals for separate meta-analytic models for each fine category of exposure. For each fine category, the first row represents the full dataset, and the second, the reduced dataset (only sample sizes of 100 or greater). The *k* column represents the numbers of correlations and the *m* column the number of unique studies. Note that all nutritional fine categories are treated as if more of the food category equalled better nutrition, and hence less adversity. Fine categories are grouped by broad category.

### Other moderators

We tested whether a series of different methodological features explained any significant amount of heterogeneity between associations (table 2; we were unable to simultaneously include exposure category and all the methodological variables in a single model for reasons of statistical power). There was no strong evidence that the study design (longitudinal vs. cross-sectional, or experimental vs correlational), the life stage of the participants (either at exposure or telomere measurement), the type of tissue, or the sex of the participants explained a significant amount of the heterogeneity, either in the full or reduced datasets.

**Table 2.** Tests of potential moderators of the association strength between exposure variables and telomere length or telomere attrition. For reasons of statistical power, potential moderators were added one at a time.

| Whole dataset |  |  |  |
| --- | --- | --- | --- |
| Moderator | Test of moderation (Q) | p-value (Q) | Parameter estimates (95% CIs) |
| Longitudinal design | Q<sub>1</sub> = 0.04 | 0.95 | Cross-sectional (ref)<br>Longitudinal -0.002 (-0.05, 0.04) |
| Experimental study | Q<sub>1</sub> = 2.84 | 0.09 | Correlational (ref)<br>Experimental -0.11 (-0.24, 0.02) |
| Life stage at exposure | Q<sub>3</sub> = 0.40 | 0.94 | Adult (ref)<br>Child 0.003 (-0.05, 0.05)<br>Embryonic 0.02 (-0.08, 0.12)<br>Not reported -0.03 (-0.16, 0.10) |
| Life stage at telomere measurement | Q<sub>3</sub> = 2.03 | 0.57 | Adult (ref)<br>Child 0.07 (-0.04, 0.18)<br>Embryonic -0.005 (-0.26, 0.25)<br>Not reported -0.08 (-0.34, 0.17) |
| Tissue type | Q<sub>2</sub> = 1.64 | 0.44 | Blood (ref)<br>Buccal cells -0.08 (-0.25, 0.10)<br>Other -0.09 (-0.26, 0.09) |
| Technique | Q<sub>3</sub> = 8.97 | 0.03* | qPCR (ref)<br>FISH -0.21 (-0.35, -0.07)<br>Southern blot -0.03 (-0.12, 0.07)<br>Telseq 0.09 (-0.30, 0.49) |
| Sex | Q<sub>2</sub> = 0.29 | 0.86 | Both sexes (ref)<br>Female -0.01 (-0.07, 0.05)<br>Male -0.01 (-0.06, 0.04) |
| Reduced dataset |  |  |  |
| Moderator | Test of moderation (Q) | p-value (Q) | Parameter estimates (95% CIs) |
| Longitudinal design | Q<sub>1</sub> = 0.00 | 0.98 | Cross-sectional (ref)<br>Longitudinal -0.0006 (-0.04, 0.04) |
| Experimental study | Q<sub>1</sub> = 3.24 | 0.07 | Correlational (ref)<br>Experimental -0.13 (-0.28, 0.01) |
| Life stage at exposure | Q<sub>3</sub> = 0.87 | 0.83 | Adult (ref)<br>Child 0.02 (-0.03, 0.06)<br>Embryonic 0.03 (-0.9, 0.15)<br>Not reported -0.01 (-0.13, 0.10) |
| Life stage at telomere measurement | Q<sub>3</sub> = 5.91 | 0.12 | Adult (ref)<br>Child 0.08 (-0.03, 0.19)<br>Embryonic 0.40 (0.005, 0.79)<br>Not reported 0.06 (-0.28, 0.39) |
| Tissue type | † | † | † |
| Technique | Q<sub>2</sub> = 0.10 | 0.95 | qPCR (ref)<br>FISH ††<br>Southern Blot -0.009 (-0.12, 0.10)<br>Telseq 0.05 (-0.30, 0.39) |
| Sex | Q<sub>2</sub> = 0.94 | 0.62 | Both sexes (ref)<br>Female 0.03 (-0.03, 0.08)<br>Male 0.004 (-0.04, 0.05) |

\* p < 0.05
† Cannot be tested because there is only a single study in any tissue type other than blood / white blood cells in the reduced dataset.
†† No FISH studies in the reduced dataset.

There was some evidence for variation in association strength by telomere measurement technique in the full dataset (table 2). Fluorescent in situ hybridization (FISH) techniques produced significantly stronger negative associations than the dominant qPCR technique. Southern blot and TelSeq associations did not differ significantly from qPCR, though TelSeq was represented in just one study. However, measurement technique was confounded with sample size in the data: FISH and Southern blot were used in relatively small-sample studies (medians 49 and 56), TelSeq in one very large study (sample size 11670), and qPCR in a range of sample sizes (median 285). We have already established that correlations were weaker in larger samples, and the order of central correlation estimates for the four techniques (FISH: *r* = −0.29, 95% CI −0.43 to −0.16; Southern blot: *r* = −0.15, 95% CI −0.20 to - 0.09; qPCR: *r* = −0.14, 95% CI −0.18 to −0.10; TelSeq: *r* = −0.04, 95% CI −0.08 to −0.01) mirrored the order of their median sample sizes. Including sample size (square-root transformed) in the model as an additional moderator, the overall moderating effect of measurement technique became marginally non-significant (Q_3_ = 7.10, p = 0.07). However, the 95% confidence interval for the parameter estimate for FISH still did not cross zero (B = −0.18, 95% CI −0.33 to −0.04, p = 0.01). Moderation by measurement technique was non-significant in the reduced dataset, though 95% of the associations in the reduced dataset used qPCR; there were no FISH associations at all, and only 15 instances of Southern blot.

## Discussion

Telomere length or attrition has been proposed as a common currency ‘psychobiomarker’ of the impact of many different types of stress and adversity on the individual. Here, we meta-analysed an exceptionally large and diverse data set consisting of 543 associations from 138 studies featuring over 400,000 human participants. The results confirm that, in the published literature, telomere length is indeed significantly associated, in the predicted direction, with a wide range of exposures including environmental hazards, smoking, psychiatric illness, psychosocial factors, socioeconomic factors, parental care, poor nutrition, and physical activity. The large heterogeneity estimate even after controlling for exposure type suggests that there is more variation in results between different studies of the same exposure than between different types of exposure.

We emphasise that because our search strategy was based on ‘stress’ and ‘adversity’, our dataset is neither exhaustive, nor a representative sample of all the work being carried out in human telomere epidemiology. We are likely to have captured almost all work on psychosocial stress, which necessarily involves one of our search terms, but only some of the studies of smoking or physical diseases. Thus, the relative abundance of exposure types in the dataset should not be interpreted as informative: for example, our dataset contains more studies of psychosocial exposures than of smoking, whereas the true abundances in the literature may well be the opposite way around [see 17,18]. However, the dataset is very large, and there are substantial numbers of studies in every broad category. Thus, it is still useful for comparing the typical strength of association of different types of exposure with telomere measures, as well as exploring cross-cutting issues. For several of the exposure variables in our dataset, there are published specialist meta-analyses covering just that exposure type. In many cases, these have appeared since we began data collection for this paper. Where such a specialist meta-analysis exists and we had more than 5 associations in our dataset, we compared the by-category results from our figure 3 with the key results of the corresponding specialist meta-analyses (table 3). There was, overall, a high degree of agreement. We view this good agreement between the specialist reviews and subsets of our dataset as confirmation that the search strategy we used yielded a sufficiently robust transect of the telomere epidemiology literature for the comparisons we have presented to be meaningful.

**Table 3.** Comparison of the present findings by fine category with key results of specialist meta-analyses, where available. Represented are central meta-analytic estimates with 95% confidence intervals. Note that we have reversed the direction of our correlations compared to figure 3 where this is necessary for the comparison.

| Exposure category | Specialist meta-analysis findings | Present findings |
| --- | --- | --- |
| Cardiovascular disease | Significant association between CVD and short TL, OR = 1.54 (1.30, 1.83) [12] | Significant negative correlation between CVD and TL, $r = -0.13$ (-0.23, -0.02) |
| Diabetes | Significant association between diabetes and short TL, OR = 1.29 (1.11, 1.50) [13] | Significant negative correlation between diabetes and TL, $r = -0.28$ (-0.41, -0.15) |
| Parkinson's disease | No significant association between TL and disease, SMD = 0.36 (-0.25, 0.96) [14] | No significant association between TL and disease, $r = 0.09$ (-0.26, 0.44) |
| Sleep apnea | Significantly shorter TL in sleep apnea, SMD = -0.03 (-0.06, -0.00) [15] | Association between TL and disease negative but not significant $r = -0.07$ (-0.27, 0.13) |
| Anxiety | Significantly shorter TL in anxiety disorders, SMD = -0.53 (-1.05, -0.01) [16] | Significantly shorter TL in anxiety disorders, $r = -0.05$ (-0.08, -0.01) |
| Depression | Significantly shorter TL in depressive disorders, SMD = -0.55 (-0.92, -0.18) [16]; $d = -0.21$ (-0.29, -0.12) [58]; $r = -0.12$ (-0.17, -0.07) [57] | Significantly shorter TL in depressive disorders, $r = -0.12$ (-0.20, -0.04), became marginally non-significant in reduced dataset |
| PTSD | Significantly shorter TL in PTSD, SMD = -1.27 (-2.12, -0.43) [16] | Significantly shorter TL in PTSD, $r = -0.17$ (-0.31, -0.03); became non-significant in reduced dataset |
| Schizophrenia | No significant association between TL and psychosis/schizophrenia, SMD = -0.2 (-0.68, 0.21) [16]; SMD = 0.34 (0.77, 154) [59]<br>Significantly shorter LTL in paranoid schizophrenia compared to controls SMD = -0.48 (-0.94, -0.03) [44] | Significant association between TL and schizophrenia, $r = -0.30$ (-0.50, -0.10); became non-significant in reduced dataset |
| Smoking | Smokers significantly shorter TL than non-smokers, SMD = -0.17 (-0.24, -0.09) [17] | Smokers significantly shorter TL than non-smokers, $r = -0.07$ (-0.11, -0.02) |
| Physical activity | Significant positive association between physical activity and TL, SMD = 0.91 (0.48, 1.35) [19] | Significant positive association between physical activity and TL, $r = 0.02$ (0.01, 0.04) |
| Stress | Weak negative correlation between perceived stress and TL, $r = -0.06$ (-0.10, -0.01); possible publication bias [18] | Weak negative correlation between perceived stress and TL, $r = -0.07$ (-0.13, -0.01); became non-significant in reduced dataset |
| Socioeconomic status other than education | No significant association with TL, SMD = 0.10 (-0.03, 0.24) [20] | No significant association with TL for composite measures, $r = 0.03$ (-0.03, 0.08) or occupation, $r = 0.00$ (-0.02, 0.03); Significant association between income and TL, $r = 0.09$ (0.01, 0.16), became non-significant in reduced dataset |
| Education | More education associated with significantly longer TL, SMD = 0.06 [0.00, 0.12] [20] | More education associated with significantly longer TL, $r = 0.05$ [0.01, 0.09] |

Abbreviations: TL: telomere length; SMD: standardized mean difference; OR: Odds ratio; $d$ : Cohen's $d$ ; $r$ : Correlation coefficient

In detecting significant negative associations between telomere variables and a wide variety of different exposures, our findings appear to support the contention that telomeres are a useful integrative ‘psychobiomarker’ [1,4]. Nonetheless, they bring to the fore a number of important caveats. The first caveat is that the observed correlations are in the range that would conventionally be considered weak or small [35]. This has methodological implications for the use of telomere measures in research. A correlation coefficient of *r* = −0.15 (our central estimate from the whole dataset) requires a sample size of 359 to be detected as significant (*p* < 0.05) with 80% power. Only 38% of the associations had this sample size. Using the central correlation estimate from the reduced dataset (*r* = −0.09), the required sample size rises to 1007 (which was met by 21% of associations). Thus, significance tests from individual small-*n* studies should not be taken as strong evidence that a given exposure variable does or does not associate with telomere length, or associates differently from other variables. Moreover, with correlations typically of small magnitude, telomere length is likely to have limited value as an indicator of adversity exposure in individual people.

The weakness of observed associations may be related to the fact that the extant literature relies almost entirely (93% of associations) on cross-sectional studies using telomere length measured at a single point in time. Where there are true environmental effects on telomere attrition, measured associations between telomere length and environmental factors in cross-sectional studies are likely to be weak, since the individual variation in telomere length at birth, which is substantially heritable, dwarfs the amount by which telomeres shorten over the life-course [36,37]. Thus, any environmental signal in cross-sectional telomere studies will be diluted by a large component of irrelevant individual variability. Longitudinal studies that examine telomere attrition, rather than telomere length, thus effectively controlling for different individual starting telomere lengths, are potentially much more powerful for detecting possible environmental influences (see [38] for discussion and [39,40] for examples of animal studies using this type of design). However, in the present dataset, we were not able to confirm that longitudinal studies produce systematically stronger negative correlations than the cross-sectional ones. This may be because longitudinal studies are few, limiting statistical power. Moreover, the follow-up tends to comprise a fairly short stretch of adult life in these human studies (mean 1576 days, s.d 882 days). A possible alternative to life-course longitudinal designs in some cases is the ‘blood-muscle’ model, where telomere length in adult muscle (where telomere length changes little) is used to estimate starting telomere length for blood [41].

The second caveat is that the literature may be affected by publication bias, a conclusion that echoes those of some narrower reviews [18]. We found evidence of stronger negative correlations in published studies with small samples, and removing the small samples nearly halved the strength of the overall association between exposure variables and telomeres. (An alternative approach to correcting for publication bias reported in the supplementary analysis document suggests an even greater degree of attenuation). Directionally stronger associations in small samples is usually taken as evidence that small studies with results contrary to prediction are being selectively withheld or rejected. Publication bias is not the only possible interpretation of this pattern, though. It could be that smaller studies measure stresses and adversities with greater precision, or use more selected participant samples so that the variation in exposure is greater, and as such genuinely detect stronger negative correlations with telomeres. Nonetheless, many of the most striking claims, for example regarding psychosocial associations with telomeres, are based on small-n findings that are atypically strong. The problem of selective appearance of associations into the literature may be worse than our findings suggest. For example, large-n epidemiological studies are likely to be published whatever the results, but authors often have degrees of freedom concerning which of many available predictor variables they report, and in how much detail. Even a slight bias towards including or providing detailed results preferentially for those measures that produce patterns conforming to expectation would suffice to distort the meta-analytic conclusions considerably.

The third caveat is that it is hard, from the present literature, to make inferences about causality in the relationships between telomere variables and exposures to stress and adversity. This is because of the overwhelming reliance on cross-sectional and correlational designs. Several of the specialist meta-analyses have concluded with calls for more longitudinal research [18,19]. It is disappointing to note that in the course of this review, we have recurrently encountered correlational findings described as if they were causal (e.g. [7,19,42,43]), and cross-sectional findings described as if they were longitudinal (e.g. [9,16,42,44]) in article titles, abstracts and discussions.

A cross-sectional correlation between telomere length and an exposure could arise for three reasons: the exposure causes telomeres to shorten (‘causality’); short telomeres cause the exposure (‘reverse causality’); or some third variable is causally related to both telomeres and the exposure (‘third variable’). Causality should not be assumed without further evidence. Reverse causality is plausible for many physical diseases. In some cases this is supported by longitudinal evidence (e.g. [41,45,46]) and Mendelian randomization studies [47,48]. Reverse causality may be possible for psychological and behavioural variables too, since short telomere length can change patterns of gene expression [49], with possible consequences for brain function. Third-variable explanations are plausible for many of the correlations described here. Childhood adversity, for example, is a third variable of potential general importance [21]. Childhood adversity is a known risk factor for a number of the variables considered here as exposures, such as poor physical and psychiatric health, smoking, and low socioeconomic status. Childhood adversity may also accelerate telomere shortening [50]. Since the highest rate of telomere shortening occurs early in life [51,52], it is perhaps more plausible that developmental conditions affect both the risk of the adult exposures and adult telomere length, than the adult exposures affecting adult telomere length directly. However, we did not find evidence in the present dataset that exposures during childhood produce significantly stronger correlations with telomere length than exposures during adulthood (though see [53] for a Cohen’s *d* effect size of −0.35 in a specialist meta-analysis of early-life adversity and telomere length).

In relation to our objective of understanding methodological sources of variation in measured associations, we were not able to reach any strong conclusions. Most of the methodological variables we recorded did not explain any significant fraction of the observed heterogeneity, but we cannot infer that they make no systematic difference. This is because many of the non-standard methodological choices in the dataset (e.g. longitudinal design, tissue other than blood, measurement technique other than qPCR) were rare. Moreover, the different features of the methodology did not vary independently of one another, or of exposure type. We found some evidence suggesting fluorescent in-situ hybridization (FISH) might produce stronger correlations with predictor variables than other measurement techniques. However, the FISH studies also featured small samples, and small sample size was associated with stronger correlations. After controlling for sample size, the moderating effect of measurement technique was attenuated. To make progress on methodological questions such as whether, for example, qPCR produces weaker associations than other techniques due to greater measurement error [23,54,55], it will be necessary to take more homogeneous sets of studies, all focussing on the same relationship, in order to isolate the consequences of this single methodological factor. For example, in recent meta-analyses of telomeres and sex [56], and telomeres and depression [57], stronger associations were found by Southern blot and/or FISH than by qPCR.

We conclude with a plea to the field. We had to exclude 125 papers because of failure to describe data in enough detail; this is nearly as many as we were able to include (138). Common omissions were simple, such as not providing means and standard deviations per group, not providing sufficient detail of regression models, or providing only a p-value for the key result. Moreover, there may have been cases where researchers measured more variables than those they reported. These failings could easily be addressed by more careful reporting of statistics, better refereeing, and, above all, fostering a culture in which all raw data are made freely available. Given the subtlety of any associations between telomere dynamics and environmental exposures, it will be necessary to pool our collective evidence in order to understand them. It is a great waste if much of that evidence is not usable for meta-analysis.

## Ethics statement

This study was based on secondary analysis of study-level data already in the public domain, and as such required no ethical approval.

## Data accessibility

Raw data and the R scripts for converting and analysing them are freely available via the Zenodo repository: http://doi.org/10.5281/zenodo.1189538.

## Authors’ contributions

G.V.P. participated in the design of the study, carried out literature searches, extracted data from the literature, designed conversion algorithms, analysed the data, and drafted the manuscript. D.N. participated in the design of the study, carried out literature searches, extracted data from the literature, analysed the data, and helped draft the manuscript. M.B. participated in the design of the study, advised on statistical analyses, and revised the manuscript for intellectual content. All authors gave approval for publication.

## Acknowledgements

We acknowledge the members of the COMSTAR lab group and other colleagues at Newcastle University for their input. We also thank Maya B. Mathur and Dan Nussey for their helpful feedback.

## Supplementary analyses for Pepper, Bateson and Nettle ‘Telomeres as integrative markers of exposure to stress and adversity: A systematic review and meta-analysis’

### 1. Background to the supplementary analyses

Certain statistical analyses used to detect outliers and publication bias cannot be straightforwardly applied to our dataset due to its multilevel structure (multiple studies each contribute multiple non-independent associations). The statistical methods required (using ‘metaplus’ and ‘weightr’ packages in R) are not currently implemented for this kind of data structure. One possibility would be to ignore the multilevel structure for these particular analyses. However, in our view this would be misleading, since most of the variation in our dataset resides at the between-study level; the multiple associations from the same study are generally highly correlated with one another (ρ = 0.87 for the simplest main analysis). Thus, to treat each association as statistically independent would be to pseudo-replicate the information from certain studies many times. In a simple meta-analytic model with no moderators, the central estimate of association is *r* = −0.15 (95% CI −0.18 to −0.11) when the multilevel structure is properly accounted for, but *r* = −0.09 (95% CI −0.10 – −0.07) if the multilevel structure is ignored and a simple random effects model is fitted. The reason for this attenuation of association strength is that studies that contribute more associations to the dataset also contribute associations that are closer to zero on average (correlation between number of associations reported and the absolute value of the mean correlation coefficient reported, *r*_136_ = - 0.27, *p* < 0.01). Thus, treating each of their data points as independent increases the influence of weak or null associations on the overall estimate.

An alternative approach that we use here is to create a ‘flat’ version of the dataset, in which one of the reported associations is chosen at random for each of the 138 studies. This produces a dataset with no multilevel structure, suitable for use with R packages ‘metaplus’ and ‘weightr’. Given that the multiple associations from the same study tend to be similar to one another, a simple random effects model of the ‘flat’ dataset leads to inferences that are broadly similar to the multilevel model of the full dataset. For example, figure S1 shows the central estimate of association between exposures and telomeres from the main multilevel model, and from ten runs of the ‘flat’ sampling procedure. The similarity suggests that, where it is not possible to use the full dataset, analyses of a ‘flat’ sample are fairly informative.

**Figure S1.**
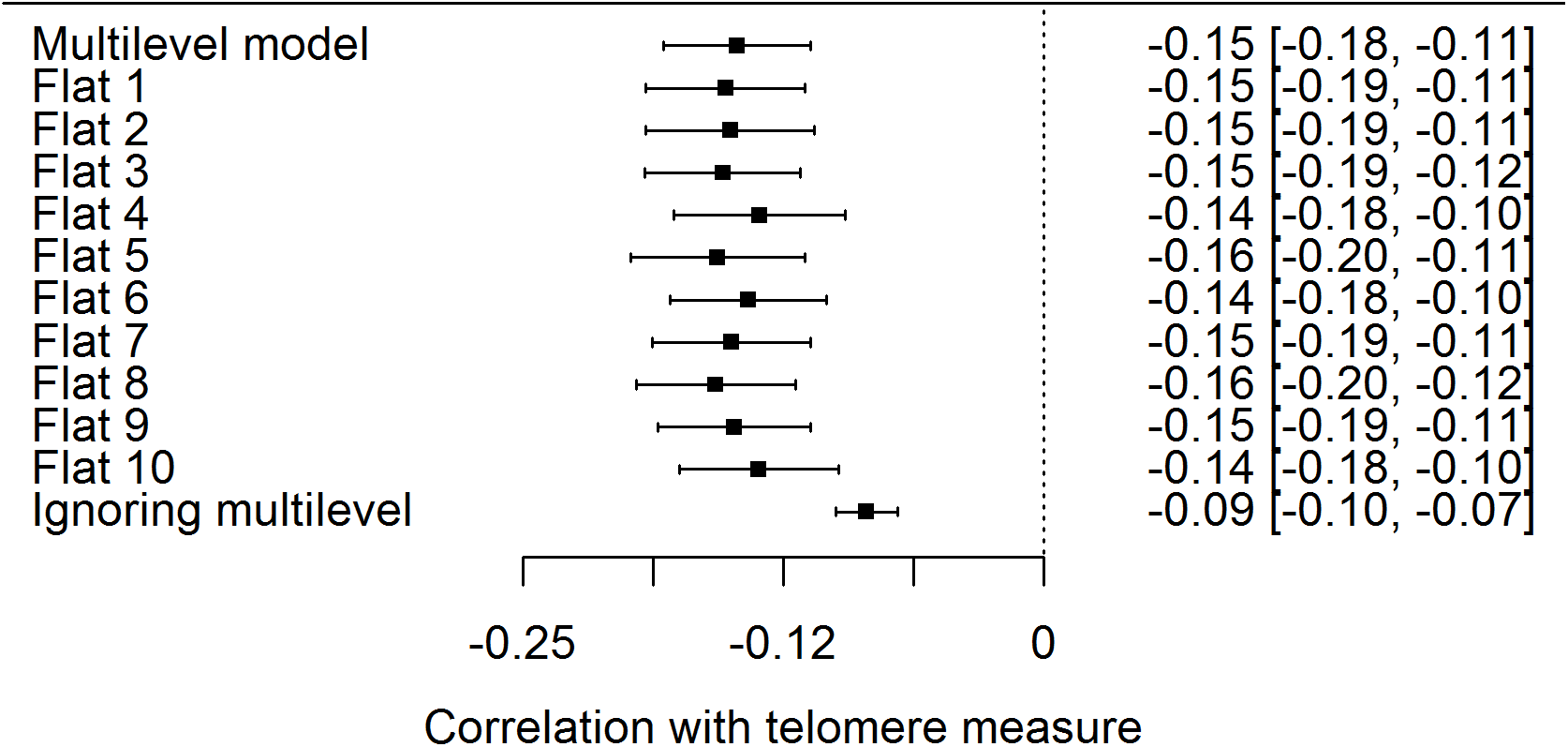
Central estimates of association between exposures and telomere measures for the main dataset analysed with a multilevel model, for ten runs of the ‘flat’ sampling procedure (see text), and for a model of the full dataset that ignores the multilevel structure and treats every association as independent.

### 2. Analysis of outliers

The standard random-effects model in meta-analysis assumes that the study-level random effects are Normally distributed. In practice, however, there may be a subset of outlier studies that depart from the central association more extremely than this. The R package ‘metaplus’ allows for the detection of the existence of such outliers. It fits a model in which the random effects are assumed to be drawn from a mixture of two distributions, one distribution for typical studies, and another, with a larger dispersion, for outlier studies (Beath, 2016). The fit of this mixture model can then be compared to the standard random-effects model: if there are studies that are marked outliers, the Bayesian Information Criterion (BIC) will be lower for the mixture model than the standard model. The studies most likely to belong to the outlier distribution can also be identified. We fit the mixture model, and a standard random-effects model, in ten ‘flat’ samples of the dataset (see section 1). Note that this procedure has limitations: a study *all* of whose associations are extreme will consistently be identified as an outlier, but a study reporting many associations only one of which is extreme will not. This should be borne in mind in what follows.

The fit for the mixture model was better than the standard model in all ten samples (mean BIC, mixture model: −24.33; standard model: −7.34). Within the mixture model, the estimated dispersion of true effects was much larger for the studies assigned to the outlier set than for the rest of the studies (mean τ, outliers: 0.35, typical: 0.07). The mean central estimate of association between exposures and telomeres for the mixture model was *r* = −0.09, somewhat weaker than the estimate from the standard model of *r* = −0.15. Thus, this analysis suggests that there is a set of outlier studies with atypically strong associations, whose aggregate effect is to make the negative correlation between exposures and telomeres appear stronger than the remaining studies would.

We examined which studies were assigned an average posterior probability of 0.9 or more of belonging to the outlier set (table S1). The 12 studies identified as outliers tended to be small-*n* (median *n* 91), and mostly reported strong negative associations between exposures and TL (10/12; the remaining two reported stronger-than-expected associations in the opposite direction). Four studies of diabetes appeared on this list, suggesting that diabetes may have genuinely stronger associations with telomere length than other categories of exposure do. Two substantial studies of sleep apnea also appeared on the list, though their findings were in opposite directions to one another, one of children suggesting longer telomere length in sleep apnea, and one in adults suggesting shorter telomere length in sleep apnea. Many of the remaining studies in table S1 had such small samples that the extreme results might simply reflect sampling variability.

**Table S1.** Studies assigned to the outlier set in the mixture-model analysis with an average posterior probability of 0.9 or greater. Note that only the first author is shown in the ‘Study’ column; for full study titles, please see processed data file in the online data archive.

| Study | Broad category | N | Comments |
| --- | --- | --- | --- |
| Adaikalakoteswari (2005) | Disease | 80 | Small study of type-2 diabetes, strong negative association of TL with disease status (patients shorter TL) |
| Castaldo (2013) | Disease | 37 | Small study on Friedrich’s ataxia, strong negative correlation of TL with duration of disease (longer duration shorter TL) |
| Defelice (2012) | Environmental hazard | 50 | Small study, very strong negative association of TL with proximity to toxic waste sites (closer to pollution shorter TL) |
| Gardner (2005) | Disease | 70 | Small longitudinal study of type-2 diabetes, strong association between change in insulin resistance and change in TL (greater rise in insulin resistance, greater TL attrition) |
| Hudson (2011) | Disease | 208 | Study of Parkinson’s disease; substantial association between white blood cell TL and disease status in opposite direction to expectation (patients longer TL) |
| Kim (2010) | Disease | 213 | Study of obstructive sleep apnea in children; substantial association between TL and sleep apnea severity in opposite direction to expectation (worse severity longer TL) |
| Krishna (2015) | Psychosocial | 33 | Small study of yoga practitioners; very large TL difference between practitioners (longer TL) and controls (shorter TL). NB. These results would be less extraordinary, though still unusually strong, if what are described as standard deviations in table 1 of the paper were actually standard errors. |
| Ma (2013) | Disease | 102 | Study of diabetes; strong negative associations between disease status and TL (patients shorter TL) |
| Marchetto (2016) | Psychosocial | 24 | Small study of prenatal stress and TL in newborn babies; strong negative association between maternal stress and TL in newborns (higher maternal stress shorter newborn TL) |
| Monickaraj (2012) | Disease | 290 | Study of type-2 diabetes, strong negative association between disease status and TL (patients shorter TL) |
| Savolainen (2014b) | Disease | 1948 | Large study of sleep apnea; substantial negative correlation between disease status and TL (patients shorter TL) |
| Snetselaar (2015) | Disease | 532 | Study of interstitial lung disease, strong negative associations between disease status and TL (patients shorter TL) |

### 3. Further analyses of publication bias

The model of Vevea and Hedges (1995) allows for assessment of, and correction for, different types of publication bias. The model simultaneously estimates the relative probability of observation of associations that lie within specified *p*-value bands (such as *p* < 0.05 versus *p* >= 0.05), as well as the overall meta-analytic association adjusted for the differential probability of observation of associations in the given bands. The model is implemented in R package ‘weightr’. We produced 10 ‘flat’ samples of our dataset (see section 1), and fitted a standard random-effects model, plus the Vevea and Hedges model, specifying the following *p*-value regions: significant association (*p* < 0.05) in the opposite direction to the hypothesis that stress and adversity shorten telomeres; non-significant trend against the hypothesis; significant trend in the direction of the hypothesis; and significant association (*p* < 0.05) in the direction of the hypothesis. The mean unadjusted association was *r* = −0.15, as expected, but the mean after adjustment for publication bias was *r* = −0.03. Thus, this analysis supports the suggestion we make in the main paper that there may be publication bias, and that it may lead to an inflated estimate of the strength of the negative association between exposures and telomere measures.

The mean relative probabilities of observation were 8.37 for a non-significant trend against the hypothesis; 21.25 for a non-significant trend in the direction of the hypothesis; and 12.94 for significant association in support of the hypothesis. These numbers are relative to a significant association contrary to the hypothesis. That is, 12.94 indicates that a significant association in the direction of the hypothesis is 12.94 more times more likely to be published, and hence appear in the dataset, than a significant association contrary to the hypothesis. The fact that all the numbers are greater than 1 suggests that every other type of result is more likely to be published than a significant finding that stress and adversity are associated with longer telomeres. In small samples, such associations should be found from time to time, because the true associations are weak, and precision of measurement is low. It is possible that researchers dismiss them as implausible or anomalous when they are found, and this explains their rarity in the dataset. This analysis suggests that the publication bias issue in this literature is not so much differential suppression of non-significant results (after all, the category with the highest relative probability of observation is actually non-significant trends in the direction of the hypothesis), but differential suppression of results that go significantly contrary to expectation.

Figure S2 visualizes this pattern. It repeats figure 1b of the main paper, but with points coloured by whether the original authors reported the associations as significant or not. As can be seen, there are plenty of non-significant associations, especially where their direction is as expected. However, there is a marked paucity of significant associations in the positive direction, even though the very broad width of the triangle where *n* is small suggests that these ought to exist.

**Figure S2.**
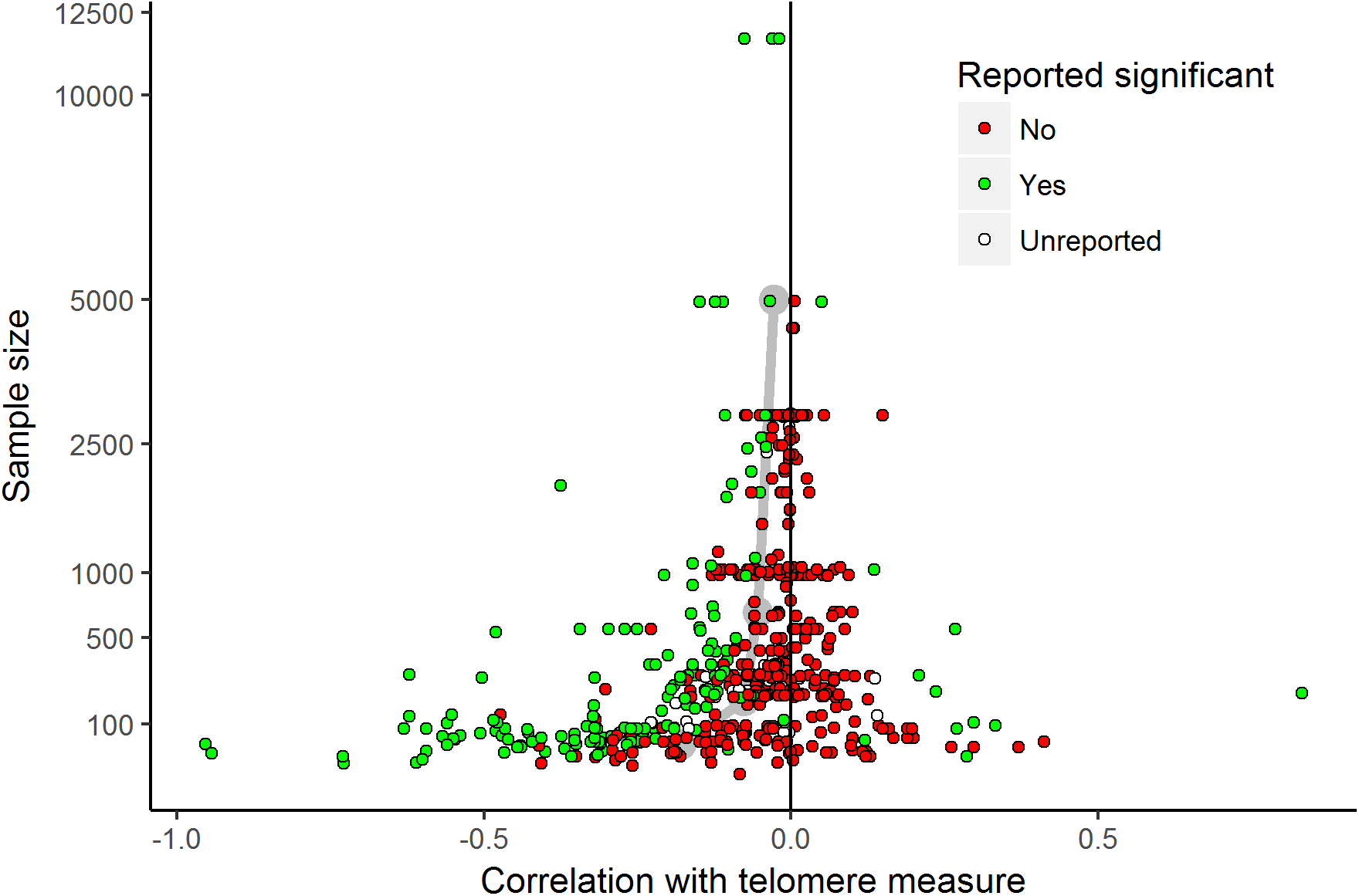
Funnel plot of sample size against observed correlation between telomere measure and exposure variable, with associations coloured to indicate whether the original authors reported them as significant or not.

